# Multiple sites on glycoprotein H (gH) functionally interact with the gB fusion protein to promote fusion during herpes simplex virus (HSV) entry

**DOI:** 10.1101/2022.12.07.519554

**Authors:** Qing Fan, Daniel P. Hippler, Yueqi Yang, Richard Longnecker, Sarah A. Connolly

## Abstract

Enveloped virus entry requires fusion of the viral envelope with a host cell membrane. Herpes simplex virus type 1 (HSV-1) entry is mediated by a set of glycoproteins that interact to trigger the viral fusion protein glycoprotein B (gB). In the current model, receptor-binding by gD signals a gH/gL heterodimer to trigger a refolding event in gB that fuses the membranes. To explore functional interactions between gB and gH/gL, we used a bacterial artificial chromosome (BAC) to generate two HSV-1 mutants that show a small plaque phenotype due to changes in gB. We passaged the viruses to select for restoration of plaque size and analyzed second-site mutations that arose in gH. HSV-1 gB was replaced either by gB from saimiriine herpesvirus type 1 (SaHV-1) or by a mutant form of HSV-1 gB with three alanine substitutions in domain V (gB3A). To shift the selective pressure away from gB, the gB3A virus was passaged in cells expressing gB3A. Sequencing of passaged viruses identified two interesting mutations in gH, including gH-H789 in domain IV and gH-S830N in the cytoplasmic tail (CT). Characterization of these gH mutations indicated they are responsible for the enhanced plaque size. Rather than being globally hyperfusogenic, both gH mutations partially rescued function of the specific gB version present during their selection. These sites may represent functional interaction sites on gH/gL for gB. gH-H789 may alter the positioning of a membrane-proximal flap in the gH ectodomain, whereas gH-S830 may contribute to an interaction between the gB and gH CTs.

**IMPORTANCE:** Enveloped viruses enter cells by fusing their envelope with the host cell membrane. Herpes simplex virus type 1 (HSV-1) entry requires the coordinated interaction of several viral glycoproteins, including gH/gL and gB. gH/gL and gB are essential for virus replication and both proteins are targets of neutralizing antibodies. gB fuses the membranes after being activated by gH/gL, but the details of how gH/gL activates gB are not known. This study examined the gH/gL-gB interaction using HSV-1 mutants that displayed reduced virus entry due to changes in gB. The mutant viruses were grown over time to select for additional mutations that could partially restore entry. Two mutations in gH (H789Y and S830N) were identified. The positions of the mutations in gH/gL may represent sites that contribute to gB activation during virus entry.

## INTRODUCTION

For most alphaherpesviruses, the core viral fusion machinery is comprised of the receptor-binding protein glycoprotein D (gD), the gH/gL heterodimer, and the fusion protein gB (1, 2). In the current model of virus entry, gD binds to one of several entry receptors. This receptor binding activates gH/gL which then triggers gB to insert into the target cell and refold, causing fusion of the viral and cellular membranes and entry of the virus particle into the cell. The entry glycoproteins are targets of neutralizing antibodies and defining how they interact could provide a basis for the development of vaccine candidates or entry inhibitors for future research or clinical use.

Multiple structures of gD, gH/gL, and gB have been determined (3-7), but the full details of the interactions among these glycoproteins during entry have not been established. Studies of neutralizing antibody epitopes have modeled sites of glycoprotein interaction (2, 4, 8, 9). Physical interactions among the entry glycoproteins also have been demonstrated (10-13), but glycoprotein complexes are difficult to capture, presumably because the glycoprotein interactions are low affinity and/or transient. gH/gL and gB are conserved among all herpesviruses, so understanding their interaction is particularly important for understanding herpesvirus entry.

The goal of this study was to use *in vitro* natural selection to identify functional interactions between gB and other entry glycoproteins. Two versions of gB with impaired fusion function were introduced into herpes simplex virus type 1 (HSV-1) and the viruses were passaged to select for secondsite mutations that rescue entry function. The first approach used a previously studied HSV-1 gB mutant and the second approach used gB from another species of alphaherpesvirus.

Approach one: Using structure-based mutagenesis, we previously identified three mutations in gB (I671A/H681A/F683A) that inhibit fusion (14). These mutations in the domain V arm of gB were designed to disrupt interactions between the arm and the domain III coil in the post-fusion form of gB. HSV-1 carrying these three mutations (gB3A virus) showed a small plaque phenotype, impaired growth, and delayed entry kinetics (15). Previously, serial passage of this gB3A virus selected for second-site mutations that enhanced plaque size (16, 17), but all of the new mutations mapped within the gB gene.

We hypothesized that we could shift the selective pressure away from gB3A by expressing gB3A in the cells used for passaging. By transiently complementing virus with cell-expressed gB3A during passage, we predicted we could select mutations in HSV-1 entry glycoproteins that restore entry by promoting an interaction with gB3A. To further shift the selective pressure away from gB, we also passaged virus deleted for gB on the gB3A-expressing cells.

Approach two: Herpesvirus entry glycoprotein interactions are often species-specific (18-21). We previously showed that gB from the saimiriine herpesvirus 1 (SaHV-1) mediated low levels of cell-cell fusion (15% of wild-type levels) when coexpressed with HSV-1 gD and gH/gL (20). SaHV-1 is a primate alphaherpesvirus with a gB homolog that has 65% sequence identity to HSV-1 gB. We hypothesized that SaHV-1 gB does not function well in fusion with HSV-1 gD and gH/gL because SaHV-1 gB interacts with these proteins poorly. In this study, we predicted that HSV-1 carrying SaHV-1 gB in place of HSV-1 gB would have reduced virus entry. We predicted that we could select for mutations in HSV-1 entry glycoproteins that promote an interaction with SaHV-1 gB by serially passaging this virus and screening for the restoration of normal entry over time.

Using both approaches, serial passage of viruses carrying impaired gB resulted in the acquisition of second-site mutations. Characterization of these mutations demonstrated that two mutations in gH (gH-H789Y and gH-S830N) partially restored gB function in fusion and plaque formation. Interestingly, rather than having a global hyperfusogenic effect, both gH mutations partially rescued the function of the specific gB version that was present during their selection. The results suggest that more than one domain of gH functionally interacts with gB during entry.

## RESULTS

### Selection of HSV-SaHVgB^pass^ viruses

Based on previous cell-cell fusion data showing that SaHV-1 gB coexpressed with HSV-1 gD, gH, and gL did not mediate high levels of fusion (20), we hypothesized that HSV-1 carrying SaHV-1 gB would show a small plaque size. To create a virus encoding SaHV-1 gB in the background of HSV-1, the SaHV-1 gB gene was recombined into a gB-null HSV-1 BAC (pQF282) to generate a new BAC (pQF397). Vero-Cre cells, which express Cre recombinase to excise the BAC backbone, were transfected with the HSV-SaHVgB BAC or wild-type (WT) BAC (pGS3217) and infectious virus was harvested. The virus encoding SaHV-1 gB was designated HSV-SaHVgB.

Vero cells were infected with the BAC-derived HSV-SaHVgB WT HSV-1 viruses at 0.01 plaque forming units (PFU)/cell. HSV-SaHVgB virus exhibited a growth defect, requiring seven days to reach full cytopathic effect (CPE), compared to only three days for WT virus. As expected, HSV-SaHVgB virus formed small plaques (Fig. 1A). Three days post-infection, plaques formed by HSV-SaHVgB virus on Vero cells were 25-fold smaller on average than WT HSV-1 plaques (Fig. 1B).

**Figure 1.**
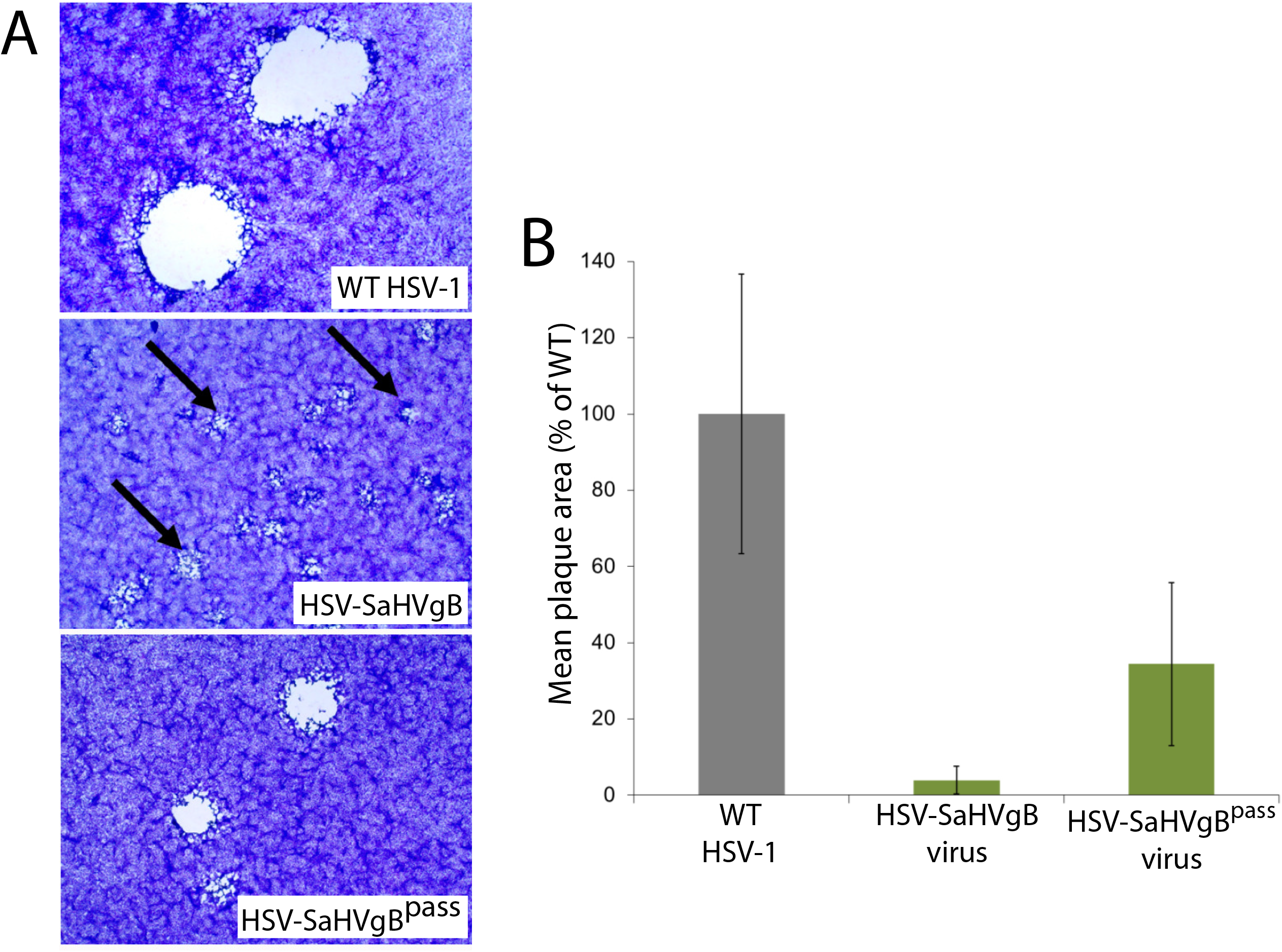
Plaque morphology of HSV-SaHVgB and HSV-SaHVgB^pass^ viruses. Vero cells were infected with WT HSV-1 (GS3217), HSV-SaHVgB (passage 1), or HSV-SaHVgB^pass^ (passage 20, lineage B) virus at an MOI of 0.01 for three days. (A) Cells were stained with Giemsa and plaques were imaged at 40X magnification. The arrows indicate small plaques. (B) The plaque sizes on Vero cells were calculated by measuring the radius of at least 50 plaques of each virus. Plaque sizes are presented as a percentage of WT plaque area. Error bars represent standard deviation.

We hypothesized that the HSV-SaHVgB virus plaque size would be increased by mutations that enhance fusion, specifically by mutations that enhance interactions between SaHV-1 gB and the HSV-1 entry glycoproteins, such as gH/gL. The locations of these mutations could reveal sites of functional importance for glycoprotein interaction and/or regulation.

To select for mutations in SaHV-1 gB or HSV-1 entry glycoproteins that could restore plaque size, HSV-SaHVgB was passaged serially in Vero cells. Four independent virus samples were passaged in parallel (Table 1). Cells were infected at a multiplicity of infection (MOI) of 0.01, virus stocks were harvested at full CPE, and stocks were titered after each passage. As the passages progressed, the time required to reach full CPE decreased and plaques size increased noticeably for all lineages. By passage 20, plaque size for one representative virus (lineage B) increased an average of 9-fold compared to the plaque size generated by a virus stock passaged only once (Fig. 1A, 1B). The passaged virus stocks were designated HSV-SaHVgB^pass^.

**Table 1.**
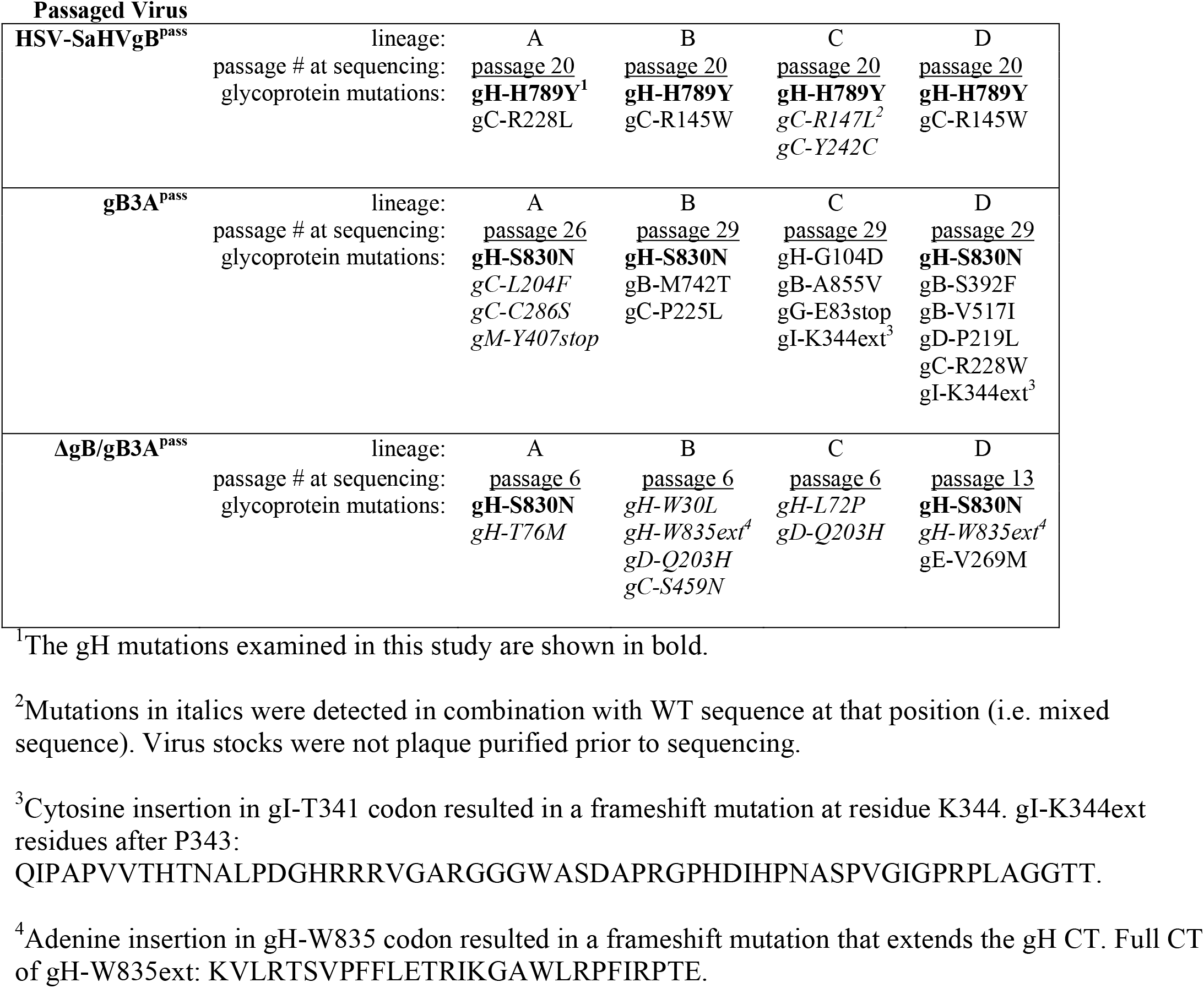
Glycoprotein mutations acquired in passaged virus isolates.

### Selection of HSV-1ΔgB/gB3A^pass^ and gB3A^pass^ viruses

In our previous work, HSV-1 encoding gB3A in place of WT gB (gB3A virus) exhibited delayed entry and a small plaque phenotype (15). The small plaque phenotype that was rescued by second-site mutations selected during serial passage; however, all of the second-site mutations occurred in gB (16, 17). We hypothesized that mutations in other entry glycoproteins also would be able to rescue the small plaque size conferred by gB3A and that those mutations would map to functional sites on the glycoproteins. To drive the selection pressure towards genes other than gB, we created an HSV-1 deleted for the gB gene that carried gB3A protein in its envelope (designated ΔgB/gB3A virus).

To produce ΔgB/gB3A virus, a Vero cell line expressing gB3A was generated and designated Vero-gB3A (Fig. 2). The gB3A gene with its native promoter was cloned into a pTuner plasmid such that gB3A expression would be induced upon HSV infection. The construct was designed to generate a bicistronic mRNA encoding gB3A and EGFP. Vero cells were transfected with this plasmid, subjected to selection with G418, and screened for GFP expression upon infection. To generate ΔgB/gB3A virus, the Vero-gB3A cells were transfected with a gB-null HSV-1 BAC (pQF282) (15), which encodes the red fluorescence protein (RFP) tdTomato gene with a nuclear localization signal under the control of a CMV promoter. Three weeks after transfection, expression of both RFP from the HSV-1 BAC and GFP from bicistronic mRNA was apparent (Fig. 2). ΔgB/gB3A virus was harvested from these cells.

**Figure 2.**
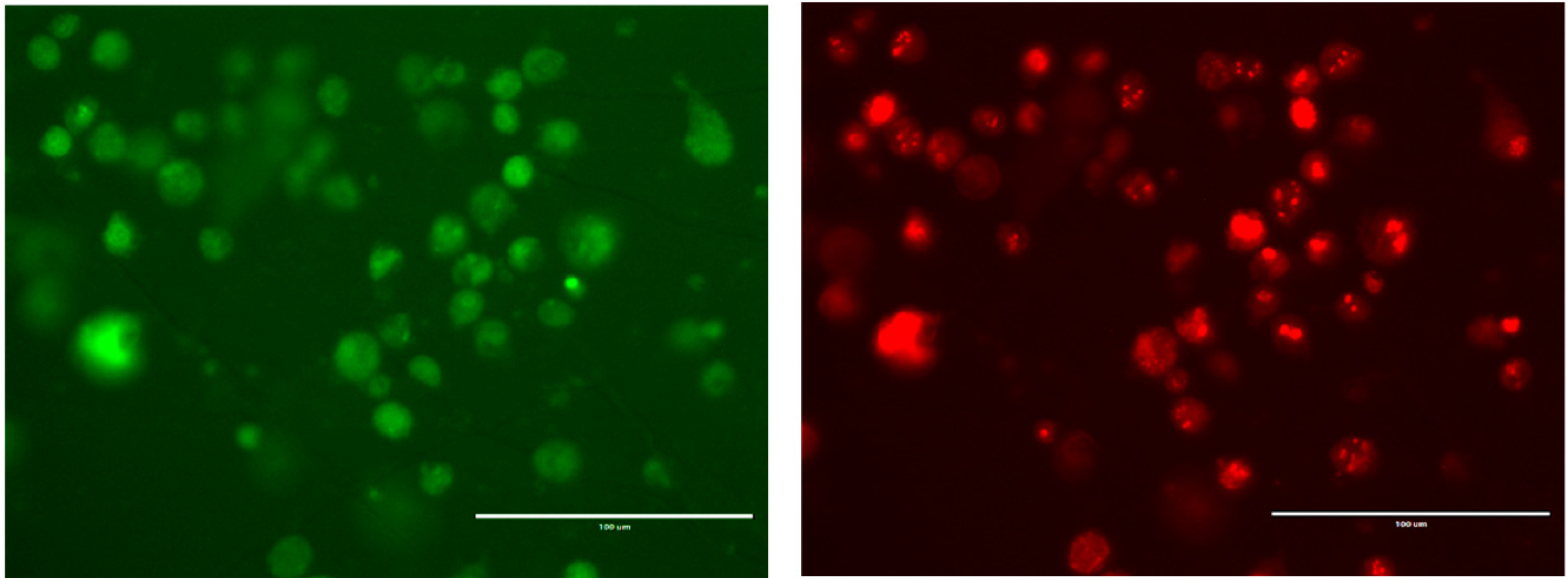
Generation of ΔgB/gB3A virus. Vero cells expressing gB3A under the native viral promoter (Vero-gB3A cells) were transfected with an HSV-1 BAC deleted for the gB gene (pQF282). After three weeks at 37°C, cells were imaged at 200X magnification for GFP (left) encoded by the gB3A expression plasmid and RFP (right) encoded by the virus. The scale bar represents 100 μm.

To select for mutations in entry glycoproteins that could restore plaque size, ΔgB/gB3A virus was passaged serially in Vero-gB3A cells. Four independent virus samples were passaged in parallel (Table 1). Cells were infected at an MOI of 0.01, supernatants were harvested at full CPE, and stocks were titered after each passage on Vero-gB3A cells. The passaged viruses were designated ΔgB/gB3A^pass^.

Using a similar logic and approach, gB3A virus (which carries the gB3A gene in place of gB) (15) also was passaged serially in Vero-gB3A cells. Four independent lineages of gB3A virus were passaged in parallel (Table 1) and the passaged viruses were designated gB3A^pass^.

Consistent with our previous study (15), gB3A virus formed small plaques on Vero cells (Fig. 3A, B). Similarly, plaques of gB3A virus on Vero-gB3A cells were small. After 26 passages, the average plaque size of gB3A^pass^ virus (lineage D) on Vero-gB3A cells had increased by nine-fold. Consistent with an enhanced growth phenotype, the time required to reach full CPE after infection decreased as the number of passages for gB3A^pass^ virus increased. Unexpectedly, the plaque size for ΔgB/gB3A^pass^ virus did not increase noticeably after passaging on Vero-gB3A cells (Fig 3A); however, the time required to reach full CPE for ΔgB/gB3A^pass^ decreased as passage number increased.

**Figure 3.**
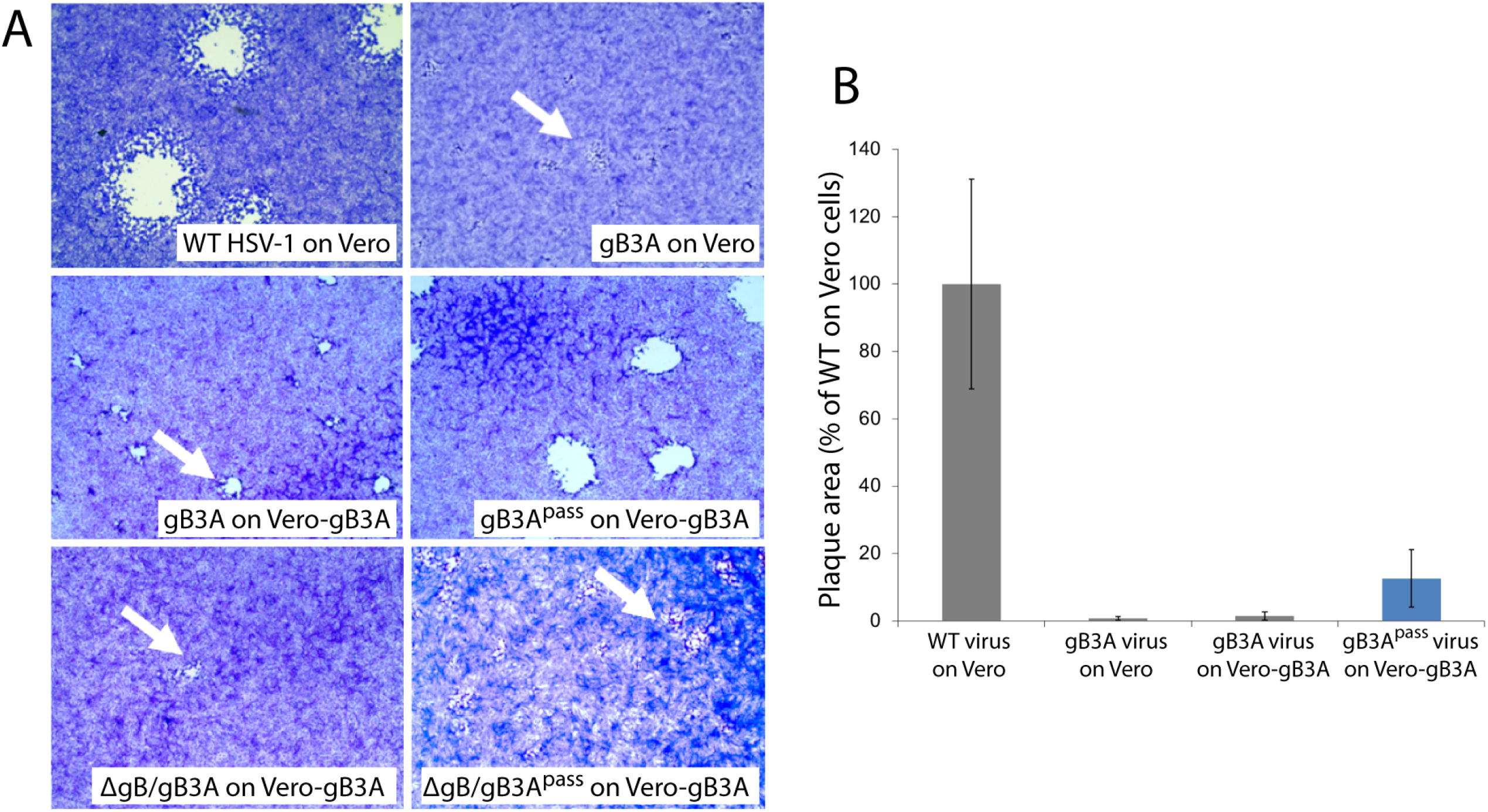
Plaque morphology of gB3A and gB3A^pass^ viruses. Plaques of WT, gB3A, gB3A^pass^ (passage 26, lineage D), ΔgB/gB3A, and ΔgB/gB3A^pass^ (passage 4, lineage D) viruses on Vero or Vero-gB3A cells are shown. Cells were infected at an MOI of 0.01 and stained with Giemsa three days post-infection. The arrows indicate small plaques. (C) Plaque sizes were calculated by measuring the radius of at least 50 plaques of WT, gB3A, and gB3A^pass^ (passage 26, lineage D) viruses on Vero or Vero-gB3A cells. Plaque sizes are presented as a percentage of WT plaque area. Error bars represent standard deviation.

### Passaged viruses have enhanced cell entry

We hypothesized that second-site mutations in the passaged viruses may have increased plaque size by rescuing an entry defect imparted by gB3A or SaHV-1 gB. To investigate an entry phenotype, we performed an entry assay using CHO cells that carry the *lacZ* gene under the control of the HSV ICP4 promoter and express the gD-receptor nectin-1 (M3A cells) or the the gD-receptor HVEM (M1A cells). Cells were infected with the original stocks of gB3A virus or HSV-SaHVgB virus and with the passaged virus stocks. Entry was assayed by measuring β-galactosidase activity (22, 23). As expected, gB3A virus and HSV-SaHVgB virus showed greatly reduced entry compared to WT virus (Fig. 4). The gB3A^pass^ virus (passage 26, lineage A) and HSV-SaHVgB^pass^ virus (passage 20, lineage B) showed enhanced entry compared to their parental strains, demonstrating a two to four-fold increase in entry.

**Figure 4.**
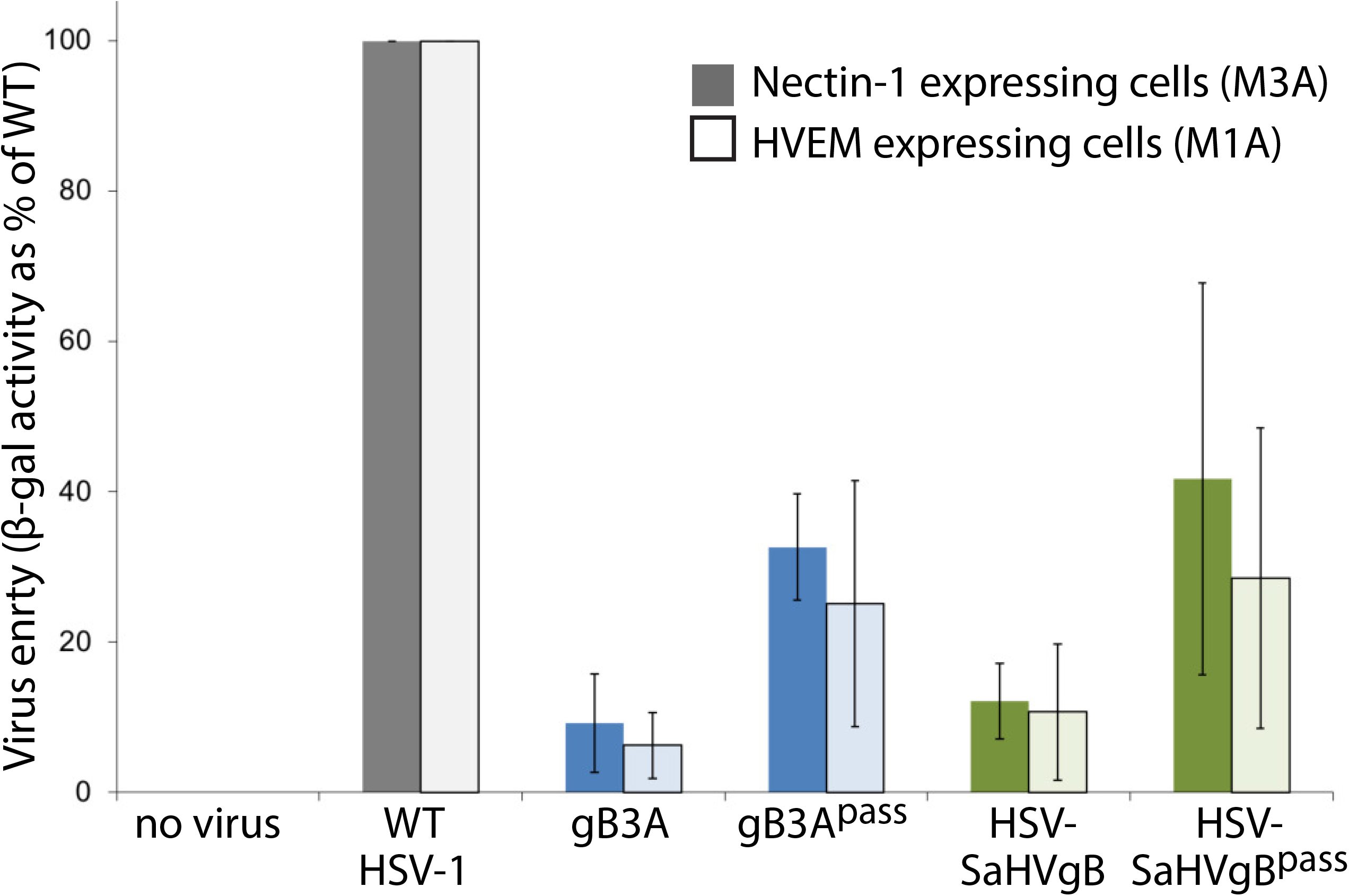
Entry of passaged viruses into cells. CHO-K1 cells that carry *lacZ* under control of the ICP4 promoter and express nectin-1 (M3A cells, dark bars) or HVEM (M1A cells, light bars) were incubated overnight with virus stocks at an MOI of 10, including gB3A^pass^ (passage 26, lineage A) and HSV-SaHVgB^pass^ (passage 20, lineage B). The β-galactosidase production was used as a measure of virus entry. Entry is shown as a percentage of WT (GS3217) virus. Error bars represent the standard deviation of three independent trials.

### Passaged viruses acquired mutations in glycoprotein H

Stocks of HSV-SaHVgB^pass^ gB3A^pass^, and ΔgB/gB3A^pass^ viruses were harvested and genomic DNA was sequenced for four independent lineages of each. For comparison, HSV-SaHVgB and gB3A virus genomes also were sequenced. For each passaged virus stock, mutations were identified in multiple glycoproteins (Table 1).

Interestingly, HSV-SaHVgB^pass^ viruses did not acquire second site mutations in SaHV-1 gB, but the gH mutation H789Y in the gH cytoplasmic tail was present in all four independent lineages. Additional mutations arose in gC as well. For the gB3A^pass^ and ΔgB/gB3A^pass^ viruses, the gH mutation S830N arose in five of the eight stocks sequenced. Additional mutations in other glycoproteins were also identified (Table 1). Two of the mutations acquired in gB, M742T and A885V, were characterized previously after they were identified by passaging gB3A virus on Vero cells (16, 17).

### Cellular expression of WT gH reduces plaque size for HSV-SaHVgB^pass^ and gB3A^pass^ viruses

To examine whether the acquired mutations in gH contributed to the increased plaque sizes for gB3A^pass^ and HSV-SaHVgB^pass^ viruses, Vero cells expressing WT gH (Vero-VgHC4 cells) were infected with the passaged viruses. We predicted that if the mutant gH proteins were promoting growth of these mutant viruses, for example through an enhanced an interaction with the mutant gB, cellular expression WT gH would compete with the mutant gH and reduce plaque size. Vero cells were used because they readily form plaques. Vero cells and Vero-VgHC4 cells were infected with equal number of PFU per well, as judged by titers on Vero cells. The plaques on Vero-VgHC4 cells were smaller on average than the plaques on Vero cells (Fig. 5). WT virus plaque sizes were similar on Vero and Vero-VgHC4 cells. These results support the hypothesis that the mutations acquired in gH contribute to the larger plaque size observed for gB3A^pass^ and HSV-SaHVgB^pass^ viruses.

**Figure 5.**
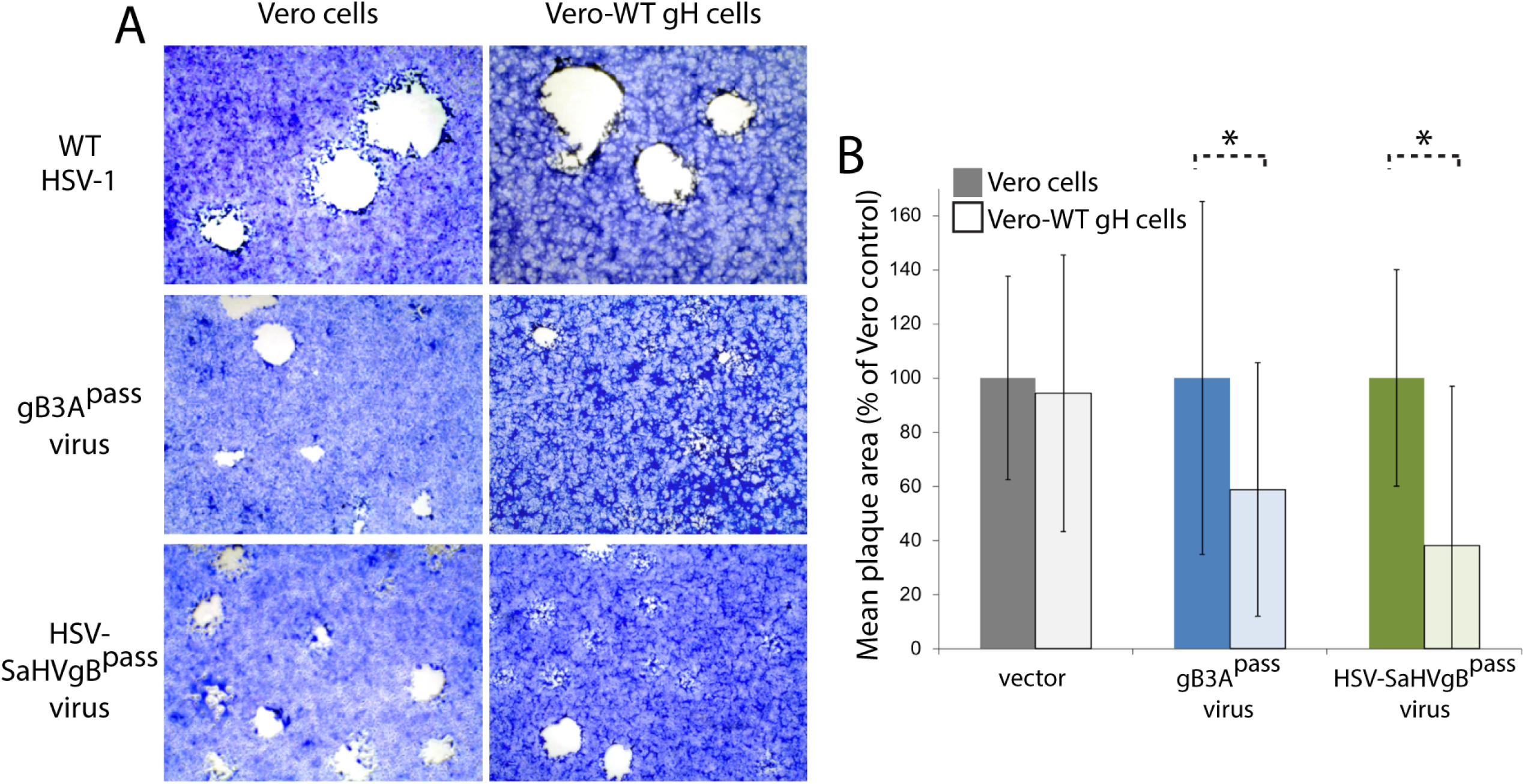
Impact of WT gH expression on growth of passaged viruses. Vero cells (dark bars) and Vero cells expressing WT gH (VgHC4 cells, light bars) were infected with WT, gB3A^pass^ (passage 26, lineage A) or HSV-SaHVgB^pass^ (passage 21, lineage B) viruses. (A) Cells were stained with Giemsa three days post-infection. (B) Plaque sizes are expressed as a percentage of plaque size on Vero cells. The error bars represent standard deviations. Asterisks indicate a significant difference in plaque size between the Vero and VgHC4 cells (Mann-Whitney U test, p≤0.01).

### Cellular expression of gH-H789Y or gH-S830N enhances plaque size for HSV-SaHVgB and gB3A viruses

We predicted that if the newly acquired gH mutations were responsible for the enhance growth observed for the passaged mutant gB viruses, cellular expression of the new gH mutants would enhance the growth of the original gB3A virus or HSV-SaHVgB virus. To examine whether specific mutations in gH contributed to the increased plaque sizes for gB3A^pass^ and HSV-SaHVgB^pass^ viruses, we expressed the gH mutants in cells and then infected with the original stocks of gB3A virus or HSV-SaHVgB virus. We focused on the gH mutations that were selected independently multiple times, gH-S830N and gH-H789Y (Fig. 1). These mutations were cloned individually into a gH expression construct. Vero cells were transfected with a plasmid encoding gH-S830N, the mutation identified in gB3A^pass^ viruses, and then infected with gB3A virus. Similarly, Vero cells were transfected with a plasmid encoding gH-H789Y, the mutation identified in HSV-SaHVgB^pass^ viruses, and then infected with HSV-SaHVgB. For comparison, Vero cells transfected with empty vector were infected in parallel. For both gB3A virus and HSV-SaHVgB virus, expression of the corresponding gH mutant increased plaque size (Fig. 6). These results are consistent with the gH mutations partially rescuing the growth defect imparted by the gB mutations.

**Figure 6.**
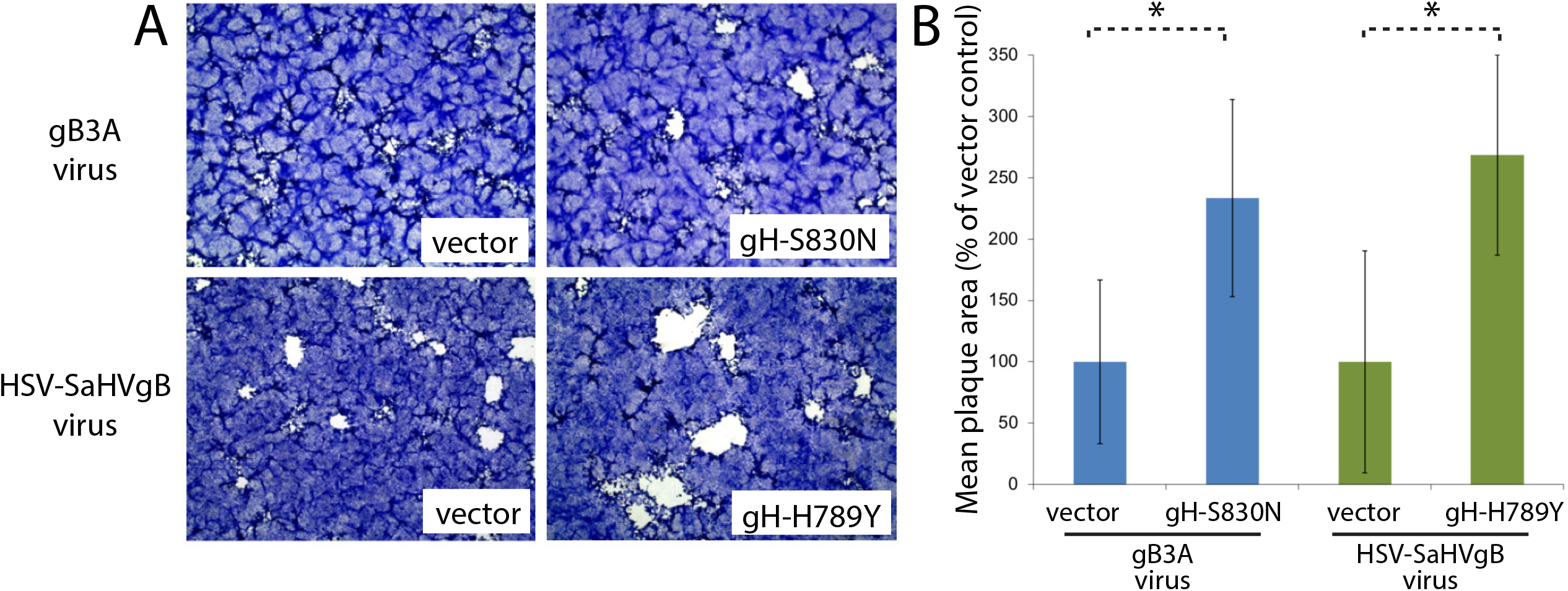
Impact of mutant gH expression on growth of mutant viruses prior to passage. Vero cells were transfected with plasmids encoding gH-S830N or gH-H789Y and then infected with gB3A or HSV-SaHVgB viruses, respectively. Cells transfected with empty vector were infected in parallel. (A) Cells were stained with Giemsa three days post-infection. (B) Plaque sizes are expressed as a percentage of plaques sizes on cells transfected with vector. The error bars represent standard deviations. Asterisks indicate a significant difference in plaque size between the Vero cells transfected with gH versus vector (Mann-Whitney U test, p≤0.01).

### gH-H789Y enhances syncytium formation mediated by SaHV-1 gB

We hypothesized that the gH mutations isolated in the gB3A^pass^ and HSV-SaHVgB^pass^ viruses were selected due to functional interactions with the gB mutants present in these viruses. We used a syncytia assay to investigate whether the gH mutants impacted the fusion mediated by WT or mutant gB. B78-H1 cells stably expressing nectin-1 (C10 cells) were transfected with the four entry glycoproteins required for fusion (gD, gB, gH, and gL), swapping in gH or gB mutants. C10 cells were used because they can form large syncytia. When co-expressed with WT gB, both gH-S830N and gH-H789Y showed reduced syncytium formation, with gH-S830N reducing the number of syncytia to below 20% of that seen with WT gH (Fig. 7A). In alignment with previous results using a cell-cell fusion assay in CHO-K1 cells (15, 20), cells expressing WT gH and either gB3A or SaHV-1 gB formed fewer syncytia than cells expressing WT gB. Co-expression of gH-S830N with gB3A did not increase the number of syncytia mediated by gB3A substantially. In contrast, co-expression of gH-H789Y with SaHV-1 gB greatly increased syncytium formation mediated by SaHV-1 gB, compared to WT gH co-expression (Fig. 7A, 7B). The finding that gH-H789Y enhances fusion mediated by SaHV-1 gB but not HSV-1 gB suggest that gH-H789Y functionally interacts with SaHV-1 gB specifically.

**Figure 7.**
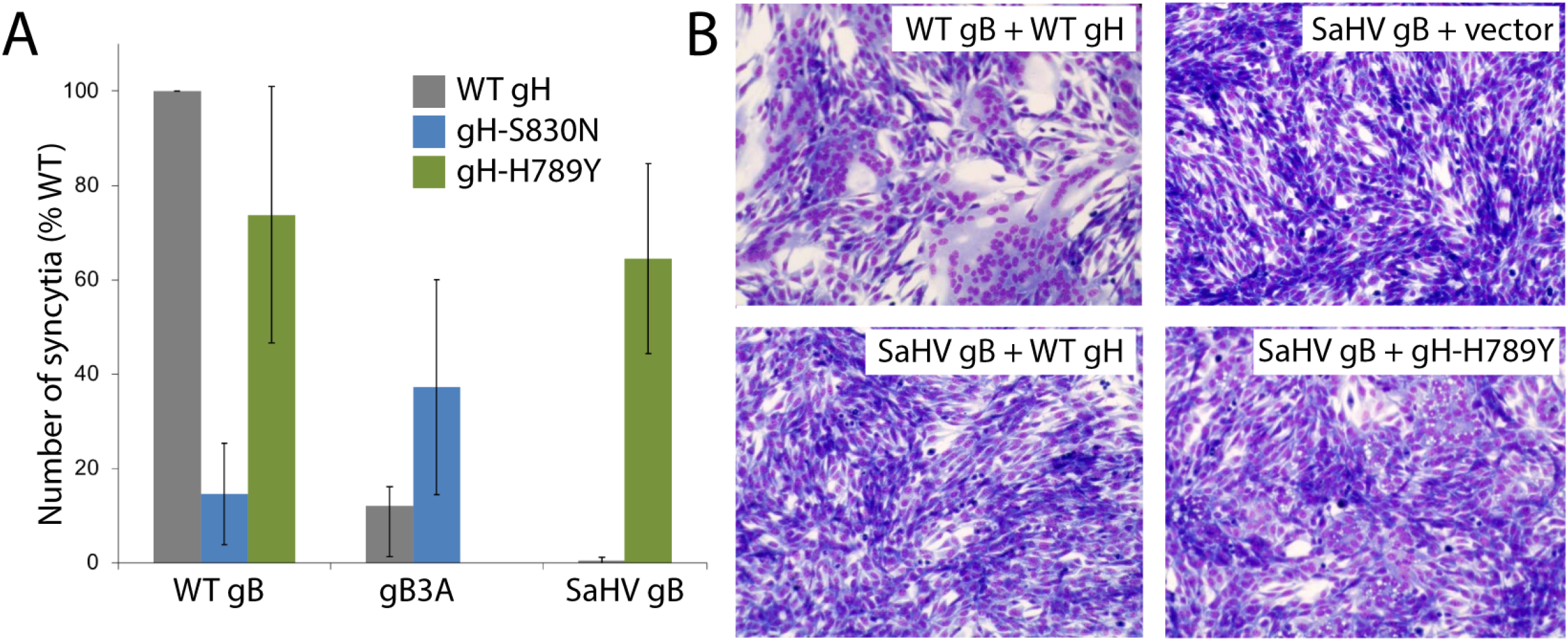
Syncytia formation upon coexpression of gH and gB mutants. (A) Cells expressing nectin-1 (B78H1 C10 cells) were transfected with plasmids encoding gD, gL, a version of gB (WT, gB3A, or SaHV-1 gB) and a version of gH (WT, gH-S830N, or gH-H789Y). Cells were stained and imaged using EVOS Cell Imaging System. Syncytia were counted and data are expressed as a percentage of the number of syncytia formed by when WT glycoproteins were expressed. The means of at least three independent determinations are shown. (B) Representative images of syncytia are shown.

### gH-H789Y and gH-S830N enhance fusion mediated by SaHV-1 gB and gB3A, respectively

To further investigate the function of these gH mutants in fusion, we used a reporter-gene cell-cell fusion assay. One set of CHO-K1 cells (effector cells) was transfected with plasmids encoding T7 polymerase, gD, gL, gB, and gH, swapping in mutant gB or gH constructs. A second set of CHO-K1 cells (target cells) was transfected with plasmids carrying the luciferase gene under the control of the T7 promoter and nectin-1 receptor. Effector and target cells were mixed and luciferase activity was measured after six hours to quantify cell-cell fusion. CHO-K1 cells were used because they provide strong signals in this assay. Cell surface expression of the WT gH and the gH mutants was determined by CELISA, using the MAb 53S on a duplicate set of effector cells.

gH-S830N and gH-H789Y were expressed at WT levels (Fig. 8A). As previously reported (15, 20), gB3A and SaHV-1 gB mediated low levels of fusion when co-expressed with WT gH. When gH-S830N or gH-H789Y was co-expressed with WT gB, fusion was reduced compared to WT gH (Fig. 8B), indicating that these mutations are not globally hyperfusogenic. In contrast, when gH-S830N was co-expressed with gB3A, fusion was enhanced compared to WT gH activity with gB3A (Fig. 8C). Similarly, when gH-H789Y was co-expressed with SaHV-1 gB, fusion was enhanced compared to WT HSV-1 gH activity with SaHV-1 gB (Fig. 8D). The enhancement observed for gH-H789Y was more pronounced than the enhancement observed for gH-S830N, consistent with the results from the syncytia assay (Fig. 7A). Taken together, the results indicate that these gH mutants specifically partially counteract the fusion deficiency imparted by gB3A or SaHV-1 gB.

**Figure 8.**
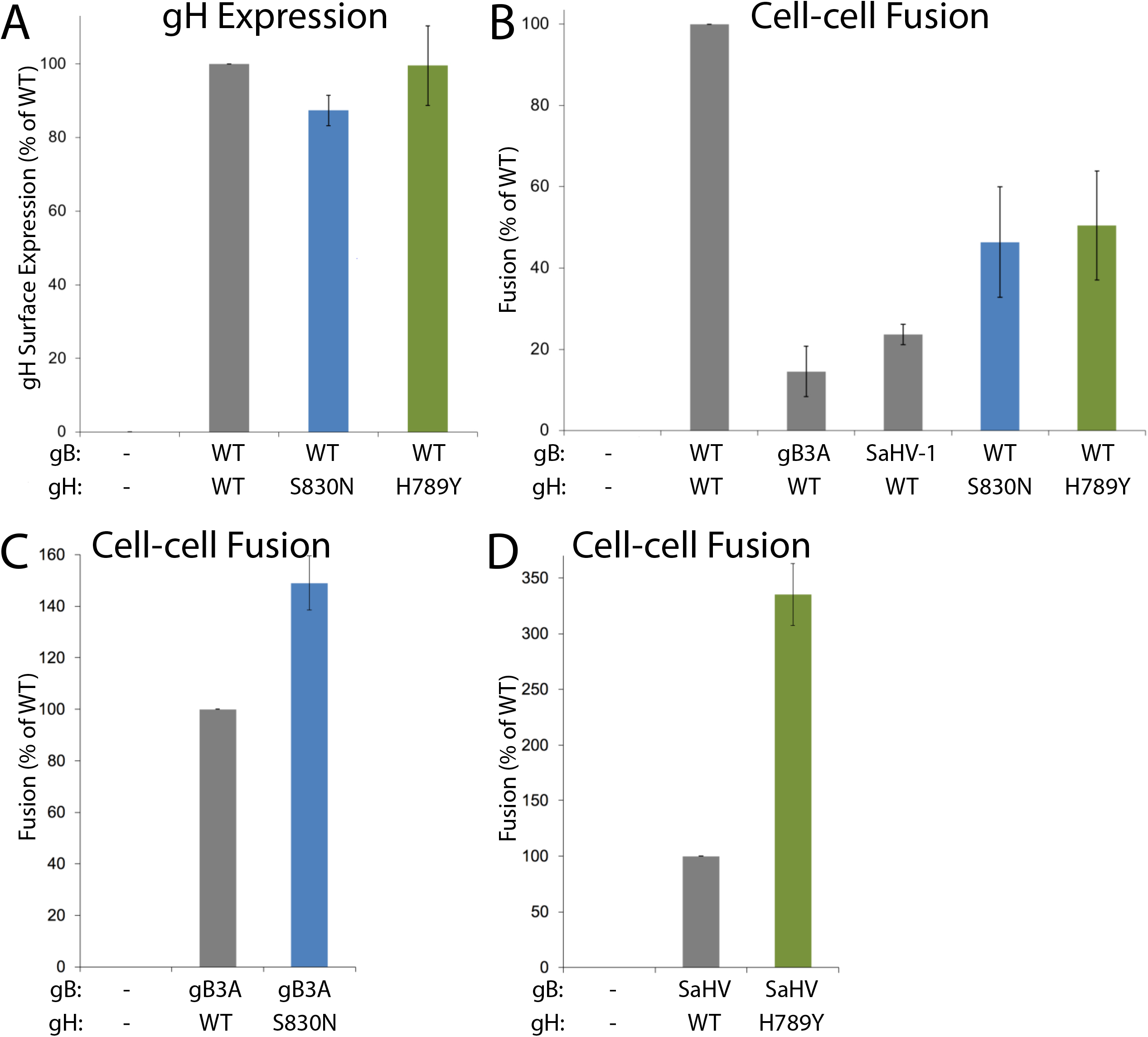
Cell-cell fusion mediated by gH and gB mutants. One set of CHO-K1 cells (effector cells) was transfected with plasmids encoding a version of gB (WT, gB3A, SaHV-1 gB, or empty vector), a version of gH (WT, gH-S830N, gH-H789Y, or empty vector), gD, gL and T7 polymerase. gB and gH versions are indicated below each graph. A second set of CHO-K1 cells (target cells) was transfected with plasmids carrying the luciferase gene under the control of the T7 promoter and nectin-1 receptor. Target and effector cells were co-cultured for six hours and luciferase activity was measured as an indication of cell-cell fusion. Cell-surface expression of gH was determined by CELISA using a duplicate set of effector cells and anti-gH MAb 53S. (A) Mutant gH surface expression when coexpressed with WT gH, gL, and gD. Data are expressed as a percentage of WT gH expression and background signals from cells transfected with vector alone are subtracted. (B) Fusion mediated by gB or gH mutants coexpressed with WT gH or WT gB. Data are expressed as a percentage of fusion in the presence of both WT gH and WT gB. (C) Fusion mediated by gB3A coexpressed with WT gH or gH-S830N. Data are expressed as a percentage of gB3A fusion in the presence of WT gH. (D) Fusion mediated by SaHV-1 gB coexpressed with WT gH or gH-H789Y. Data are expressed as a percentage of SaHV-1 gB fusion in the presence of WT gH. For all graphs, background fusion signals detected after transfection with vector instead of glycoproteins were subtracted from the values.

## DISCUSSION

Herpesvirus entry into cells is complex because it requires the concerted effort of multiple viral glycoprotein, as opposed to most enveloped viruses that require only one or two viral glycoproteins for entry. Mapping the physical and functional glycoprotein interaction sites has been challenging because glycoprotein complexes are difficult to capture. This study used structure-based mutagenesis and *in vitro* natural selection to examine how mutations impact glycoprotein interactions. Building on previous studies, we introduced functionally-impaired gB mutants into the HSV genome and use serial passage to select for second site mutations that rescue entry function. In the case of the gB3A virus, to shift the selective pressure away from gB and explicitly select for mutations outside of gB, we passaged virus on cells expressing gB3A. A similar approach was used previously to isolate extragenic suppressors of an HSV-1 UL34 mutant (24, 25).

Passage of these viruses selected for novel gH mutations, multiple times independently. Despite being conserved in all herpesviruses, the role of the HSV gH/gL heterodimer in entry is defined incompletely. The current model of entry places gH/gL between gD and gB in the sequence of events, with receptor-binding by gD initiating a change in gH/gL that triggers gB to refold to drive fusion. Interactions of all of the glycoprotein combinations (gD-gH/gL, gD-gB, and gH/gL-gB) have been demonstrated using a split-luciferase assay (13) and bimolecular fluorescence complementation (BiFC) assays (10, 12).

Purified forms of the gD and gH/gL ectodomains have been shown to interact directly using surface plasmon resonance (SPR) (8, 11). Anti-gH/gL MAbs that block the gD-gH/gL interaction detected by SPR map to the flexible N-terminus of gH (residues 19-47) and the C-terminus of gL (resides 48, 55, 77) (Fig. 9), suggesting that gD binds to the gH/gL N-terminus (8). Studies of panels of neutralizing anti-gH/gL and anti-gD antibodies support this location for gD binding (9). Chimeric constructs of HSV gH and SaHV-1 gH also mapped a species-specific gD functional interaction to the N-terminal half of the gH/gL ectodomain (26).

**Figure 9.**
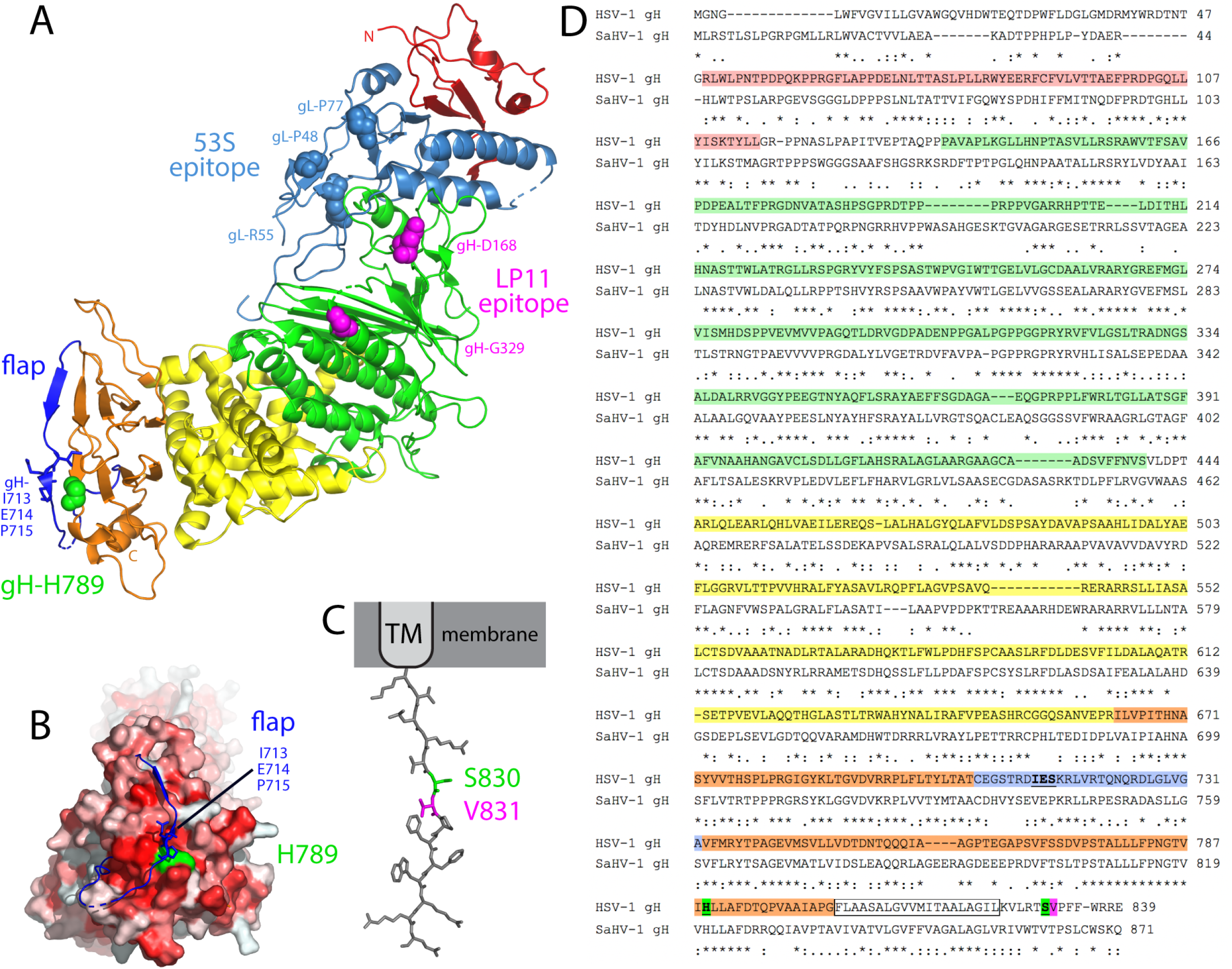
Structural model of gH/gL. (A) The boot-shaped crystal structure of HSV-2 gH/gL (PDB ID 3M1C) is presented from a side-view, with the gH N- and C-termini labeled. gL (light blue) associates with domain I (red) and domain II (green) of gH in the membrane-distal N-terminal half of the complex. Domain III (yellow) and domain IV (orange) comprise the membrane-proximal half of the heterodimer with the TM anchor following domain IV (unresolved). Point mutations in gL (light blue spheres) reduce the binding of the neutralizing MAb 53S, an antibody that blocks a gD-gH/gL interaction detected by SPR (8). These residues mark a putative gD-binding site on gH/gL. Point mutations in gH (magenta spheres) reduce the binding of the neutralizing MAb LP11, an antibody that reduces a gB-gH/gL interaction detected by BiFC (4). These residues mark a potential gB-binding site on gH/gL. The base of domain IV is capped by a conserved flap (dark blue) comprised of residues C706-A732 (36). Residue gH-H789 (green spheres) lies directly beneath this flap, contacting gH flap residues 713-715 (blue sticks). (B) The gH/gL structure has been rotated approximately 45 degrees, providing a view of the “toe” of the boot. A surface rendering of gH generated in the absence of the flap (blue) is shown. Residues are colored according to hydrophobicity (i.e. darker red is more hydrophobic). Residue H789 (green) lies beneath the flap and contacts residues 713-715 (blue sticks). Movement of the flap would expose a hydrophobic patch (red) near H789. (C) The 14 residue CT of gH has been modeled in PyMol as an extended strand. Residue S830 (green) neighbors V831 (magenta), a residue that is proposed to interact with a pocket in the gB CT (47). Images were generated using PyMol (https://pymol.org/2/). (D) An amino acid alignment of HSV-1 gH (GenBank accession number AFE62849) and SaHV-1 gH (GenBank accession number ADO13806.1) was generated using MAFFT version 7. Identical (asterisks) and similar (colons or periods) residues are marked and amino acid numbers listed. gH domains are colored as in A. The TM is boxed. gH-H789 and gH-S830 are green and gH-V831 is magenta. The domain IV flap is blue with residues predicted to contact H789 underlined. Residue 715 is proline in the HSV-2 gH/gL structure.

The gB interaction site on gH/gL is less well defined than the gD interaction site on gH/gL. Purified forms of the HSV gH/gL and gB ectodomains were shown to associate at low pH using a co-flotation liposome binding assay (27). Similarly, human cytomegalovirus gH/gL and gB were coimmunoprecipitated and this association did not require the gB cytoplasmic domain (28). Using BiFC, the anti-gH/gL neutralizing MAb LP11 was shown to reduce the interaction between gH/gL and gB (4), suggesting that gB binds to the gH/gL ectodomain at a site distinct from the gD-interaction site (Fig. 9). In addition, chimeric constructs of HSV-1 gH and pseudorabies gH mapped a species-specific gB functional interaction to the membrane-proximal domain of the gH/gL ectodomain (21).

Although the gB and gH/gL ectodomains appear to interact and a purified form of the HSV gH/gL ectodomain was sufficient to trigger low levels of cell-cell fusion (29), the gH transmembrane (TM) and/or cytoplasmic tail (CT) also contribute to fusion. Mutations in the TM or CT of membrane-anchored gH inhibit fusion function without impairing protein folding (30-35). Insertion of five residues in the gH CT, immediately after the TM domain, completely abrogates fusion despite normal levels of expression (31). Removal of the last nine residues of the gH tail (i.e. S830 and beyond) results in a small plaque phenotype, reminiscent of the gB3A plaque phenotype (32, 34).

Recently, a gH/gL-gB interaction was shown using a split-luciferase assay and chimeric forms of gH comprised of combinations of HSV and Epstein Barr virus (EBV) segments (13). The gH/gL ectodomain, CT and TM each were shown to promote the gH/gL-gB physical interaction independently suggesting that gH/gL may interact with gB at multiple sites. Consistent with a model of gB interacting with multiple sites on gH/gL, the two gH mutations in the current study that partially restored virus entry in the presence of gB mutants map to different domains of gH/gL: the membrane-proximal domain of gH (H789Y) and the gH cytoplasmic tail (S830N) (Fig. 9).

Passage of HSV-1 carrying SaHV-1 gB resulted in the selection of gH-H789Y in all four independent lineages (Table 1). We concluded that the gH-H789Y mutation contributed to the larger plaque size (Fig. 1) and enhanced entry (Fig. 4) observed for the HSV-SaHVgB^pass^ viruses because (i) gH-H789Y enhanced cell-cell fusion when coexpressed with SaHV-1 gB (Fig. 7, 8), (ii) providing gH-H789Y *in trans* during infection with HSV-SaHVgB virus increased plaque size (Fig. 6), and (iii) complementing the passaged HSV-SaHVgB virus with WT gH reduced plaque size (Fig. 5).

gH-H789 is conserved in HSV-1 and SaHV-1 gH homologs, near the center of stretch of 20 amino acids in domain IV that are nearly identical for the two viruses (Fig. 9D). H789 is only 16 residues upstream of the TM domain, placing this residue close to the membrane. Based on the HSV-2 gH/gL structure, H789 lies directly beneath a previously identified “flap” that runs across the bottom edge of domain IV (36, 37) (Fig. 9A). This flap (residues C706-G732) covers a hydrophobic patch on domain IV and H789 contacts residues 713-715 in the center of the flap (Fig. 9B). Mutating H789 to tyrosine, a bulky polar residue, may disrupt interactions with the flap, causing a shift in the flap position and a conformation of gH that more readily exposes the hydrophobic patch beneath the flap. None of the gH homologs examined encode a tyrosine at this position (36). Notably, gH-H789Y does not enhance fusion when coexpressed with WT HSV gB (Fig. 8B), so the alteration to gH enhances fusion via a specific interaction with SaHV gB. These results suggest a species-specific gH-gB interaction may be enhanced by this change to the base of domain IV.

Passage of HSV-1 carrying virally-encoded gB3A or cell-expressed gB3A resulted in the selection of gH-S830N in five independent lineages (Table 1). We concluded that the gH-S830N mutation contributed to enhanced entry using the same reasoning as for the gH-H789Y mutation above. gH-S830 maps to the short gH CT (residues 825-838) that is essential for WT levels of fusion (1, 2, 38).

Two non-exclusive possibilities may explain how gH-S830N restores entry. The gH CT may influence the gH ectodomain conformation through inside-out signaling and/or the gH CT may interact directly with the gB CT. Truncations in the EBV gH CT were previously shown to reduce binding of gp42 to the gH/gL ectodomain, demonstrating that the gH CT influences the gH ectodomain structure (39). In addition, the CT of fusion proteins from multiple virus families have been shown affect fusion protein function (40-42). The gB CT regulates gB activity, potentially serving as a ‘clamp’ that stabilizes gB (5). Mutations or truncations in the gB CT enhance fusion (43-46), potentially by disrupting the clamp. During virus entry, the gH CT may act as a ‘wedge’ to disrupt the gB CT and promote fusion (5). Using mutagenesis and structural modeling, a recent study proposed that gH-V831, a residue neighboring gH-S830, may insert into a pocket in the gB CT to activate gB by disrupting interprotomer contacts and releasing the ‘clamp’. This gB pocket was identified because mutations near this site can have hyperfusogenic or hypofusogenic effects. When the gH tail was modeled as an extended strand, gH-V831 was positioned at the same distance from the membrane as the functional gB pocket.

gH-S830N may promote gB3A virus entry by enhancing an interaction of the gH CT with the gB3A CT. gH-S830N displayed enhanced fusion when coexpressed with gB3A (Fig. 8C) but not with WT gB (Fig. 8B), indicating that S830N is not globally hyperfusogenic and suggesting that the CT of gB3A may be altered compared the CT of WT gB. Multiple studies have demonstrated that that the gB CT influences activation of the gB ectodomain. The result from the current study suggests that the gB ectodomain conformation (i.e. gB3A) may also influence the gB CT conformation. The gH-S830N mutation replaces a polar residue with a non-polar residue and may facilitate gH-V831 insertion into the gB3A CT pocket by altering interactions with residue near the gB3A CT pocket. Similar to our results for gH-S830N, gH-S830A was shown recently to reduce fusion moderately when coexpressed with WT gB (47).

Additional novel mutations in glycoproteins were identified in the passaged viruses (Table 1). One of the ΔgB/gB3A^pass^ lineages acquired a mutation that extends the gH cytoplasmic tail (gH-W835ext), replacing the final three residues with nine residues. Previous work has shown that the gH tail residues past V831 also contribute to fusion. gH truncation mutants show that gH fusion activity positively correlates with gH tail length (35). Likewise, the mutation V831A reduces fusion more substantially in the context of a tail truncated after residue 832 rather than a full-length tail (30), suggesting the distal region of the tail can compensate for the V831A substitution (47). In addition, all four HSV-SaHVgB^pass^ lineages acquired mutations in gC. gC contributes to entry by binding to cell surface glycosaminoglycans (48). Recent work has shown that gC can regulate entry via a low-pH pathway and impact gB conformational changes (49, 50). Whether these gC and gH mutations contributed to the enhanced entry observed in these passaged viruses may be the subject of future work.

gB and gH/gL represent fusion machinery that is conserved across all herpesviruses, underscoring the importance of defining how gH/gL interacts with gB and triggers the conformational change in gB that mediates fusion. The current work supports the model that gH/gL regulates gB fusion function and predicts that gH/gL-gB interactions occur at multiple sites. The approach employed provides a means to genetically map interaction sites on gH/gL and potentially other proteins that have weak or transient interactions to dissect a stepwise understanding of the herpesvirus fusion mechanism.

## MATERIALS AND METHODS

### Cells and antibodies

African green monkey kidney Vero cells (American Type Culture Collection [ATCC], USA) were grown in Dulbecco modified Eagle medium (DMEM) supplemented with 10% fetal bovine serum (FBS) (ThermoFisher Scientific, USA), penicillin, and streptomycin. Vero-CRE cells were derived from Vero cells to express Cre recombinase, (kindly provided by Dr. Gregory Smith at Northwestern University) and Vero-VgHC4 cells (31) were derived from Vero cells to express WT gH. Both of the Vero-derived cell lines were grown in the same medium as Vero cells. Chinese hamster ovary (CHO-K1) (ATCC, USA) cells were grown in Ham’s F12 medium supplemented with 10% FBS. M1A and M3A cells (22) were derived from CHO-IEβ8 cells to stably express human HVEM and nectin-1, respectively. CHO-IEβ8 is a CHO-K1-derived cell line that expresses β-galactosidase under the control of an immediate early promoter. M1A and M3A cells were grown in Ham’s F-12 medium supplemented with 10% FCS, 150 μg of puromycin/mL, and 250 μg of G418/mL. C10 cells were derived from B78-H1 mouse melanoma to stably express human nectin-1 (51). They were grown in DMEM supplemented with 10% FBS and 500 μg of G418/mL. gH antibody anti-gH monoclonal antibody (MAb) 53S (52) recognizes HSV-1 gH/gL.

### Plasmids and Bacterial Artificial Chromosomes (BACs)

Plasmids and BACs generated for this study include: pQF439 (pSG5-gB), pQF440 (pSG5-gD), pQF441 (pSG5-gH), pQF442 (pSG5-gL), pQF444 (pSG5-gH-S830N), pQF445 (pSG5-gH-H789Y), pQF356 (pTuner-gB3A), pQF397 (HSV-SaHV1gB BAC), and pQF395 (an intermediate construct for BAC construction).

Plasmids encoding HSV-1 KOS strain gB (pPEP98), gD (pPEP99), gH (pPEP100) and gL (pPEP101) were previously described (53). To clone HSV-1 gB, gD, and gL into the pSG5 vector, pPEP98, pPEP99, and pPEP101 were digested with EcoRI and BglII and the fragments containing the glycoproteins were ligated into pSG5 after digestion with EcoRI and BglII. This generated plasmids pSG5-gB (pQF439), pSG5-gD (pQF440) and pSG5-gL (pQF442). To clone HSV-1 gH into the pSG5 vector, pPEP100 was digested with SacI and the fragment was blunted with T4 DNA polymerase and then cut with BglII. pSG5 was digested with EcoRI, blunted with T4 DNA polymerase, and then cut with BglII. The gH fragment was ligated to the digested pSG5 to generate pSG5-gH (pQF441).

Site-directed mutagenesis of pQF441 was used to generate pSG5-gH-S830N (pQF444) using primers 5’GGTTCTCCGGACAAATGTCCCGTTTTTTTG3’ and 5’CAAAAAAACGGGACA TTTGTCCGGAGAACC3’ and to generate pSG5-gH-H789Y (pQF445) using primers 5’CAAACGGAACCGTCATTTATTTGCTAGCCTTTGAC3’ and 5’GTCAAAGGCTAGCAAAT AAATGACGGTTCCGTTTG3’.

gB3A was expressed using the previously described plasmid pSG5-HSVgB-I671A/H681A/F683A (14). SaHV-1 gB was expressed using the previously described pCAGGS plasmid pQF77 (20).

pT7EMCLuc plasmid encoding a firefly luciferase reporter gene under the control of the T7 promoter and pCAGT7 plasmid encoding T7 RNA polymerase (53, 54) were used in the fusion assay.

To generate an inducible expression construct to make a Vero-gB3A cell line, gB3A was inserted into pTuner-IRES2 using the plasmid pUL34-IRES2-EGFP (24). This plasmid was kindly provided by Dr. Richard Roller at University of Iowa. In pUL34-IRES2-EGFP, the CMV promoter was replaced by the UL34 promoter and UL34 expression is induced by infection. We replaced UL34 with gB3A and the gB promoter such that the resulting plasmid (pQF356) encodes gB3A under control of its native promoter with bicistronic expression of EGFP. To amplify the gB3A gene with its native promoter, we used the previously described HSV-1 gB3A BAC (pQF297) (15).

pGS1439, a plasmid containing the kanamycin resistance gene (kan^R^), was used for the BAC construction and was kindly provided by Dr. Gregory Smith at Northwestern University. gB3A virus and ΔgB/gB3A viruses were generated using the previously described HSV-1 gB3A BAC (pQF297) and gB-null BAC (pQF282), respectively (15). HSV-SaHV1gB virus was generated by creating a new BAC (pQF397), using an intermediate gB-kan^R^ construct (pQF395), as described below.

### Construction of HSV-SaHVgB BAC

The HSV-SaHVgB BAC (pQF397) was generated using the gB-null BAC pQF282 (15). pQF282 was generated previously by deleting UL27 (the gB gene) from the BAC GS3217 (55). GS3217 is an HSV-1 F strain BAC that carries the red fluorescence protein (RFP) tdTomato reporter gene with a nuclear localization signal under the control of a CMV immediate early promoter. The CMV>NLS-tdTomato>pA cassette is inserted in the US5 (gJ) gene.

The HSV-SaHVgB BAC was generated using a two-step red-mediated recombination strategy (56). First, the kan^R^ gene was PCR-amplified from pGS1439 using the primers 5’-ACGGGTC**TGTACA**ACGACCGCGCCCCGGTTCCAT CGAAGAGATCACGGACGTGATCAAC GCCAA<u>AGGATGACGACGATAAGTAGGGA</u>-3’ and 5’-GAACCGGGGCGCGGTCGT**TGTACA**<u>CAACCAATTAACCAATTCTGAT-</u>3’. These primers consist of 5’ SaHV-1 gB sequence, including a BsrGI restriction site in bold, and 3’ kan^R^ homology (underlined). This PCR product was digested with BsrGI and ligated into BsrGI-digested pQF77 (pCAGGS-SaHV-1 gB) to generate pQF395. The SaHV-1 gB gene containing a kan^R^ insert was PCR-amplified from pQF395 using primers 5’-GTCCTCCAGCAC CTCGCCCCCAGGCTACCTGACGGGGGGCACGACGGGCCCCCGTAGTCCCGCCATGGCGCCTCCGGCCGCCAAGAGC-3’ and 5’-AACAAACCAAAAGATGCACATGCGGTTTAACAC CCGTGGTTTTTATTTACAACAAACCCCCCGCTACACAGCAGCGTCGTCTTCGTCC-3’. Using two-step red-mediated recombination, this PCR product was recombined into the gB-null BAC pQF282. Then the kan^R^ cassette was recombined out to generate the BAC HSV-SaHVgB (pQF397) that carries the SaHV-1 gB gene in place of HSV-1 gB. The intermediate BAC with kan^R^ insertion and the final BAC constructs were confirmed by at least four restriction enzyme digestions.

### Generating virus stocks

gB3A virus and WT HSV-1 (GS3217) were generated from BACs previously (15). To generate HSV-SaHVgB virus, the HSV-SaHVgB BAC (pQF397) was transfected into four wells of Vero-Cre cells using Lipofectamine 2000. The transfected cells were harvested 2-3 weeks after transfection using three rounds of freeze-thaw. Four independent transfections were used to generate the four lineages. To generate ΔgB/gB3A virus, four wells of Vero-gB3A cells were transfected with the gB-null BAC (pQF282) using Lipofectamine 2000 (Invitrogen, Carlsbad, CA). For lineage D, a plasmid expressing Cre recombinase was cotransfected. Virus was harvested from the cells when CPE was apparent at 2-3 weeks after transfection using three rounds of freeze-thaw. Four independent transfections were used to generate the four lineages used for passage.

### Generating Vero-gB3A cell line

Vero cells in 6-well plates were transfected with 1.5 μg/well pQF356 using Lipofectamine 2000 (Invitrogen, Carlsbad, CA). Then the cells were seeded in 100 mm plates in medium containing 800 μg/mL G418 to allow colonies to develop for two to three weeks. Approximately 100 individual colonies were seeded into 96-well plates. When the cells reached near confluence, they were transferred to 24-well plates. As these cells reached confluency, duplicate cultures were seeded in 24-plates. One set of cells was saved and one was assessed for GFP expression after an overnight infection with WT HSV-1 at an MOI of 5. Cells that screened positive for GFP expression were subjected to two additional rounds of screening. Fluorescence microscopy of was performed using an EVOS Cell Imaging Systems (AMG, Fisher Scientific).

### Determining plaque size

Plaques were visualized three days post-infection using Giemsa staining and imaged with transmitted light microscopy using EVOS Cell Imaging Systems. The average radius of randomly-selected plaques was calculated from two independent measurements. At least 50 plaques were measured for each virus sample. Using the average radius, the plaque area and the ratio of plaque size between WT and mutant viruses was determined, as described previously (16, 17).

### Entry of mutant viruses into M1A and M3A cells

As previously described (22), M1A or M3A cells growing in 96-well plates were infected at an MOI of 1. After 5 hours at 37°C, the cells were washed with phosphate buffered saline (PBS) and lysed in 50 μL/well DMEM containing 0.5% NP-40. β-galactosidase activity was measured by adding 50 μL/well of a 4.8 mg/mL solution of chlorophenol red-β-D-galactopyranoside (CPRG; Boehringer Mannheim) and reading the absorbance at 560 nm on a plate reader (Perkin Elmer). This assay was repeated at least three times.

### Virus genome sequencing and analysis

DNA was purified for sequencing from cells infected in six well plates using a blood and cell culture DNA mini kit (Qiagen). Samples were sequenced at the Northwestern University Genomics Core Facility using NextGen Illumina HiSeq SR500 sequencing. Unipro UGene (57) was used to create a trimmed reference sequence composed of the HSV-1 F strain genome (GenBank accession number GU734771.1) with the long and short terminal repeat regions deleted (58). The gB (UL27) gene in the reference sequence was replaced with either the SaHV-1 gB sequence or the KOS strain gB sequence (GenBank accession number KT899744.1) with the gB3A substitutions added. Using Geneious 8.1.9, sequencing reads were aligned to the reference genome and variants were identified. Variants present in both the revertant viruses and the original BACs were ignored, as were variants in the gJ gene due to the RFP insertion. Mutations identified in other glycoproteins are reported in Table 1. The SaHV-1 gB sequence of HSV-SaHVgB^pass^ viruses was confirmed using standard single pass DNA sequencing after PCR amplification.

### Infection of gH-expressing cells

Vero and VgHC4 (cells expressing WT gH) seeded in 6-well plates were infected at 150 pfu/well with WT, gB3A^pass^, or HSV-SaHVgB^pass^ virus. Alternatively, Vero cells seeded in 6-well plates were transfected with 1.5 μg/well of plasmids encoding gH-S830N, gH-H789Y, or vector using 5 μL Lipofectamine 2000. After an overnight incubation, cells were infected at 150 pfu/well with gB3A or HSV-SaHVgB virus. Viral titers had been determined previously on Vero cells. Cells were stained with Giemsa three days post-infection to visualize plaques.

### Syncytium formation assay

The syncytium formation assay was performed as described previously (23). C10 cells in 24-well plates were transfected with 250 ng/well each of plasmids encoding of gB, gD, gH, and gL using 2.5 μL/well FuGENE 6 (Promega, USA) in 500 μL/well DMEM with 10% FBS. When indicated, WT versions were substituted with gB and/or gH mutants. After an overnight incubation, cells were fixed with methanol and stained with Giemsa for 30 minutes. Syncytia, defined as multinucleated cells with three or more nuclei, were counted using microscopy.

### Cell-cell fusion assay

The fusion assay was performed as previously described (53). Briefly, CHO-K1 cells were seeded in 6-well plates overnight. One set of cells (effector cells) were transfected with 400 ng each of plasmids encoding T7 RNA polymerase, gB, gD, gL, and gH, using 5 μl of Lipofectamine 2000 (Invitrogen, USA). When indicated, WT versions were substituted with gB and/or gH mutants. WT glycoproteins were expressed from pSG5 plasmids, rather than pCAGGS plasmids as described previously. A second set of cells (target cells) was transfected with 400 ng of a plasmid encoding the firefly luciferase gene under control of the T7 promoter and 1.5 μg of a plasmid expressing nectin-1, using 5 μL of Lipofectamine 2000. After overnight transfection, the cells were detached with versene and resuspended in 1.5 mL/well of F12 medium supplemented with 10% FBS. Effector and target cells were mixed in a 1:1 ratio and re-plated in 96-well plates for six hours. Luciferase activity was quantified using a luciferase reporter assay system (Promega) and a Wallac-Victor luminometer (Perkin Elmer).

### Cell-based ELISA (CELISA)

To evaluate the cell surface expression of the gH mutants, CELISA staining was performed as previously described (59). During the cell-cell fusion assay, a duplicate set of CHO-K1 effectors cells was generated. After an overnight incubation, these cells were rinsed with PBS and the primary anti-gH/gL MAb 53S was added at 1:5000 dilution. After a one-hour incubation, the cells were rinsed, fixed in formaldehyde-glutaraldehyde, and incubated with biotinylated goat anti-mouse IgG (Sigma), followed by streptavidin-HRP (GE Healthcare) and HRP substrate (BioFX).

### Statistical analysis

Statistical comparison of the plaque areas was performed with a Mann-Whitney U test using SPSS (version 25). Analyses were performed using IBM SPSS statistics version 25 for Windows (IBM Corp., Armonk, NY) and SAS 9.4 (SAS Institute, Cary, NC).

## ACKNOWLEDGMENTS

We thank Richard Roller for providing the pUL34-IRES2-EGFP plasmid. We thank Gregory Smith for providing the BAC GS3217 as well as Yasushi Kawaguchi for providing the parental BAC. We thank Kevin Bohannon for generating the BAC GS3217. We thank Yi Li at IQVIA for statistical analysis. We thank Sarah Kopp and Nanette Susmarski for excellent technical assistance and members of the Longnecker laboratory for their help in these studies. Sequencing services were performed at the Northwestern University Genomics Core Facility. R.L. is the Dan and Bertha Spear Research Professor in Microbiology-Immunology. This work was supported by NIH grant AI148478.

## REFERENCES

1. Connolly SA, Jardetzky TS, Longnecker R. 2021. The structural basis of herpesvirus entry. Nat Rev Microbiol 19:110–121.

2. Gonzalez-Del Pino GL, Heldwein EE. 2022. Well Put Together-A Guide to Accessorizing with the Herpesvirus gH/gL Complexes. Viruses 14.

3. Carfi A, Willis SH, Whitbeck JC, Krummenacher C, Cohen GH, Eisenberg RJ, Wiley DC. 2001. Herpes simplex virus glycoprotein D bound to the human receptor HveA. Mol Cell 8:169–179.

4. Chowdary TK, Cairns TM, Atanasiu D, Cohen GH, Eisenberg RJ, Heldwein EE. 2010. Crystal structure of the conserved herpesvirus fusion regulator complex gH-gL. Nat Struct Mol Biol 17:882–888.

5. Cooper RS, Georgieva ER, Borbat PP, Freed JH, Heldwein EE. 2018. Structural basis for membrane anchoring and fusion regulation of the herpes simplex virus fusogen gB. Nat Struct Mol Biol 25:416–424.

6. Krummenacher C, Supekar VM, Whitbeck JC, Lazear E, Connolly SA, Eisenberg RJ, Cohen GH, Wiley DC, Carfi A. 2005. Structure of unliganded HSV gD reveals a mechanism for receptor-mediated activation of virus entry. Embo J 24:4144–4153.

7. Heldwein EE, Lou H, Bender FC, Cohen GH, Eisenberg RJ, Harrison SC. 2006. Crystal structure of glycoprotein B from herpes simplex virus 1. Science 313:217–220.

8. Cairns TM, Atanasiu D, Saw WT, Lou H, Whitbeck JC, Ditto NT, Bruun B, Browne H, Bennett L, Wu C, Krummenacher C, Brooks BD, Eisenberg RJ, Cohen GH. 2020. Localization of the Interaction Site of Herpes Simplex Virus Glycoprotein D (gD) on the Membrane Fusion Regulator, gH/gL. J Virol 94.

9. Atanasiu D, Saw WT, Cairns TM, Eisenberg RJ, Cohen GH. 2021. Using Split Luciferase Assay and anti-HSV Glycoprotein Monoclonal Antibodies to Predict a Functional Binding Site Between gD and gH/gL. J Virol doi:10.1128/JVI.00053-21.

10. Atanasiu D, Whitbeck JC, Cairns TM, Reilly B, Cohen GH, Eisenberg RJ. 2007. Bimolecular complementation reveals that glycoproteins gB and gH/gL of herpes simplex virus interact with each other during cell fusion. Proc Natl Acad Sci U S A 104:18718–18723.

11. Cairns TM, Ditto NT, Atanasiu D, Lou H, Brooks BD, Saw WT, Eisenberg RJ, Cohen GH. 2019. Surface Plasmon Resonance Reveals Direct Binding of Herpes Simplex Virus Glycoproteins gH/gL to gD and Locates a gH/gL Binding Site on gD. J Virol 93.

12. Avitabile E, Forghieri C, Campadelli-Fiume G. 2007. Complexes between herpes simplex virus glycoproteins gD, gB, and gH detected in cells by complementation of split enhanced green fluorescent protein. J Virol 81:11532–11537.

13. Pataki Z, Rebolledo Viveros A, Heldwein EE. 2022. Herpes Simplex Virus 1 Entry Glycoproteins Form Complexes before and during Membrane Fusion. mBio 13:e0203922.

14. Connolly SA, Longnecker R. 2012. Residues within the C-terminal arm of the herpes simplex virus 1 glycoprotein B ectodomain contribute to its refolding during the fusion step of virus entry. J Virol 86:6386–6393.

15. Fan Q, Kopp SJ, Connolly SA, Longnecker R. 2017. Structure-Based Mutations in the Herpes Simplex Virus 1 Glycoprotein B Ectodomain Arm Impart a Slow-Entry Phenotype. MBio 8.

16. Fan Q, Kopp SJ, Byskosh NC, Connolly SA, Longnecker R. 2018. Natural Selection of Glycoprotein B Mutations That Rescue the Small-Plaque Phenotype of a Fusion-Impaired Herpes Simplex Virus Mutant. MBio 9.

17. Fan Q, Longnecker R, Connolly SA. 2021. Herpes Simplex Virus Glycoprotein B Mutations Define Structural Sites in Domain I, the Membrane Proximal Region, and the Cytodomain That Regulate Entry. J Virol 95:e0105021.

18. Zago A, Jogger CR, Spear PG. 2004. Use of herpes simplex virus and pseudorabies virus chimeric glycoprotein D molecules to identify regions critical for membrane fusion. Proc Natl Acad Sci U S A 101:17498–17503.

19. Plate AE, Smajlovic J, Jardetzky TS, Longnecker R. 2009. Functional analysis of glycoprotein L (gL) from rhesus lymphocrytovirus in Epstein-Barr virus-mediated cell fusion indicates a direct role of gL in gB-induced membrane fusion. J Virol 83:7678–7689.

20. Fan Q, Longnecker R, Connolly SA. 2014. Substitution of herpes simplex virus 1 entry glycoproteins with those of saimiriine herpesvirus 1 reveals a gD-gH/gL functional interaction and a region within the gD profusion domain that is critical for fusion. J Virol 88:6470–6482.

21. Bohm SW, Backovic M, Klupp BG, Rey FA, Mettenleiter TC, Fuchs W. 2015. Functional Characterization of Glycoprotein H Chimeras Composed of Conserved Domains of the Pseudorabies Virus and Herpes Simplex Virus 1 Homologs. J Virol 90:421–432.

22. Krummenacher C, Rux AH, Whitbeck JC, Ponce-de-Leon M, Lou H, Baribaud I, Hou W, Zou C, Geraghty RJ, Spear PG, Eisenberg RJ, Cohen GH. 1999. The first immunoglobulin-like domain of HveC is sufficient to bind herpes simplex virus gD with full affinity, while the third domain is involved in oligomerization of HveC. J Virol 73:8127–8137.

23. Connolly SA, Landsburg DJ, Carfi A, Whitbeck JC, Zuo Y, Wiley DC, Cohen GH, Eisenberg RJ. 2005. Potential nectin-1 binding site on herpes simplex virus glycoprotein D. J Virol 79:1282–1295.

24. Roller RJ, Haugo AC, Kopping NJ. 2011. Intragenic and extragenic suppression of a mutation in herpes simplex virus 1 UL34 that affects both nuclear envelope targeting and membrane budding. J Virol 85:11615–11625.

25. Vu A, Poyzer C, Roller R. 2016. Extragenic Suppression of a Mutation in Herpes Simplex Virus 1 UL34 That Affects Lamina Disruption and Nuclear Egress. J Virol 90:10738–10751.

26. Fan Q, Longnecker R, Connolly SA. 2015. A Functional Interaction between Herpes Simplex Virus 1 Glycoprotein gH/gL Domains I and II and gD Is Defined by Using Alphaherpesvirus gH and gL Chimeras. J Virol 89:7159–7169.

27. Cairns TM, Whitbeck JC, Lou H, Heldwein EE, Chowdary TK, Eisenberg RJ, Cohen GH. 2011. Capturing the herpes simplex virus core fusion complex (gB-gH/gL) in an acidic environment. J Virol 85:6175–6184.

28. Vanarsdall AL, Howard PW, Wisner TW, Johnson DC. 2016. Human Cytomegalovirus gH/gL Forms a Stable Complex with the Fusion Protein gB in Virions. PLoS Pathog 12:e1005564.

29. Atanasiu D, Saw WT, Cohen GH, Eisenberg RJ. 2010. Cascade of events governing cell-cell fusion induced by herpes simplex virus glycoproteins gD, gH/gL, and gB. J Virol 84:12292–12299.

30. Silverman JL, Heldwein EE. 2013. Mutations in the cytoplasmic tail of herpes simplex virus 1 gH reduce the fusogenicity of gB in transfected cells. J Virol 87:10139–10147.

31. Jackson JO, Lin E, Spear PG, Longnecker R. 2010. Insertion mutations in herpes simplex virus 1 glycoprotein H reduce cell surface expression, slow the rate of cell fusion, or abrogate functions in cell fusion and viral entry. J Virol 84:2038–2046.

32. Browne HM, Bruun BC, Minson AC. 1996. Characterization of herpes simplex virus type 1 recombinants with mutations in the cytoplasmic tail of glycoprotein H. J Gen Virol 77:2569–2573.

33. Harman A, Browne H, Minson T. 2002. The transmembrane domain and cytoplasmic tail of herpes simplex virus type 1 glycoprotein H play a role in membrane fusion. J Virol 76:10708–10716.

34. Wilson DW, Davis-Poynter N, Minson AC. 1994. Mutations in the cytoplasmic tail of herpes simplex virus glycoprotein H suppress cell fusion by a syncytial strain. J Virol 68:6985–6993.

35. Rogalin HB, Heldwein EE. 2015. Interplay between the Herpes Simplex Virus 1 gB Cytodomain and the gH Cytotail during Cell-Cell Fusion. J Virol 89:12262–12272.

36. Backovic M, Dubois RM, Cockburn JJ, Sharff AJ, Vaney MC, Granzow H, Klupp BG, Bricogne G, Mettenleiter TC, Rey FA. 2010. Structure of a core fragment of glycoprotein H from pseudorabies virus in complex with antibody. Proc Natl Acad Sci U S A 107:22635–22640.

37. Fuchs W, Backovic M, Klupp BG, Rey FA, Mettenleiter TC. 2012. Structure-based mutational analysis of the highly conserved domain IV of glycoprotein H of pseudorabies virus. J Virol 86:8002–8013.

38. Cairns TM, Connolly SA. 2021. Entry of Alphaherpesviruses. Curr Issues Mol Biol 41:63–124.

39. Chen J, Jardetzky TS, Longnecker R. 2016. The Cytoplasmic Tail Domain of Epstein-Barr Virus gH Regulates Membrane Fusion Activity through Altering gH Binding to gp42 and Epithelial Cell Attachment. MBio 7.

40. Janaka SK, Gregory DA, Johnson MC. 2013. Retrovirus glycoprotein functionality requires proper alignment of the ectodomain and the membrane-proximal cytoplasmic tail. J Virol 87:12805–12813.

41. Chen J, Zhang X, Jardetzky TS, Longnecker R. 2014. The Epstein-Barr virus (EBV) glycoprotein B cytoplasmic C-terminal tail domain regulates the energy requirement for EBV-induced membrane fusion. J Virol 88:11686–11695.

42. Waning DL, Russell CJ, Jardetzky TS, Lamb RA. 2004. Activation of a paramyxovirus fusion protein is modulated by inside-out signaling from the cytoplasmic tail. Proc Natl Acad Sci U S A 101:9217–9222.

43. Gage PJ, Levine M, Glorioso JC. 1993. Syncytium-inducing mutations localize to two discrete regions within the cytoplasmic domain of herpes simplex virus type 1 glycoprotein B. J Virol 67:2191–2201.

44. Baghian A, Huang L, Newman S, Jayachandra S, Kousoulas KG. 1993. Truncation of the carboxy-terminal 28 amino acids of glycoprotein B specified by herpes simplex virus type 1 mutant amb1511-7 causes extensive cell fusion. J Virol 67:2396–2401.

45. Fan Z, Grantham ML, Smith MS, Anderson ES, Cardelli JA, Muggeridge MI. 2002. Truncation of herpes simplex virus type 2 glycoproein B increases its cell surface expression and activity in cell-cell fusion, but these properties are unrelated. JVirol 76:9271–9283.

46. Chowdary TK, Heldwein EE. 2010. Syncytial phenotype of C-terminally truncated herpes simplex virus type 1 gB is associated with diminished membrane interactions. J Virol 84:4923–4935.

47. Pataki Z, Sanders EK, Heldwein EE. 2022. A surface pocket in the cytoplasmic domain of the herpes simplex virus fusogen gB controls membrane fusion. PLoS Pathog 18:e1010435.

48. Shukla D, Spear PG. 2001. Herpesviruses and heparan sulfate: an intimate relationship in aid of viral entry. J Clin Invest 108:503–510.

49. Sari TK, Gianopulos KA, Nicola AV. 2020. Conformational Change in Herpes Simplex Virus Entry Glycoproteins Detected by Dot Blot. Methods Mol Biol 2060:319–326.

50. Gianopulos KA, Komala Sari T, Weed DJ, Pritchard SM, Nicola AV. 2022. Conformational Changes in Herpes Simplex Virus Glycoprotein C. J Virol 96:e0016322.

51. Miller CG, Krummenacher C, Eisenberg RJ, Cohen GH, Fraser NW. 2001. Development of a syngenic murine B16 cell line-derived melanoma susceptible to destruction by neuroattenuated HSV-1. Mol Ther 3:160–168.

52. Showalter SD, Zweig M, Hampar B. 1981. Monoclonal antibodies to herpes simplex virus type 1 proteins, including the immediate-early protein ICP 4. Infect Immun 34:684–692.

53. Pertel PE, Fridberg A, Parish ML, Spear PG. 2001. Cell fusion induced by herpes simplex virus glycoproteins gB, gD, and gH-gL requires a gD receptor but not necessarily heparan sulfate. Virology 279:313–324.

54. Okuma K, Nakamura M, Nakano S, Niho Y, Matsuura Y. 1999. Host range of human T-cell leukemia virus type I analyzed by a cell fusion-dependent reporter gene activation assay. Virology 254:235–244.

55. Stults AM, Smith GA. 2019. The Herpes Simplex Virus 1 Deamidase Enhances Propagation but Is Dispensable for Retrograde Axonal Transport into the Nervous System. J Virol 93.

56. Tischer BK, Smith GA, Osterrieder N. 2010. En passant mutagenesis: a two step markerless red recombination system. Methods Mol Biol 634:421–430.

57. Okonechnikov K, Golosova O, Fursov M, team U. 2012. Unipro UGENE: a unified bioinformatics toolkit. Bioinformatics 28:1166–1167.

58. Szpara ML, Gatherer D, Ochoa A, Greenbaum B, Dolan A, Bowden RJ, Enquist LW, Legendre M, Davison AJ. 2014. Evolution and diversity in human herpes simplex virus genomes. J Virol 88:1209–1227.

59. Lin E, Spear PG. 2007. Random linker-insertion mutagenesis to identify functional domains of herpes simplex virus type 1 glycoprotein B. Proc Natl Acad Sci U S A 104:13140–13145.

